# pyrpipe: a python package for RNA-Seq workflows

**DOI:** 10.1101/2020.03.04.925818

**Authors:** Urminder Singh, Jing Li, Arun Seetharam, Eve Syrkin Wurtele

## Abstract

The availability of terabytes of RNA-Seq data and continuous emergence of new analysis tools, enable unprecedented biological insight. However, implementing RNA-Seq analysis pipelines in a reproducible, flexible manner is challenging as data gets bigger and more complex. Thus, there is a pressing requirement for frameworks that allows for fast, efficient, easy-to-manage, and reproducibile analysis. Simple scripting has many challenges and drawbacks. We have developed a python package, python RNA-Seq Pipeliner (**`pyrpipe`**) that enables straightforward development of flexible, reproducible and easy-to-debug computational pipelines purely in python, in an object-oriented manner. **`pyrpipe`** provides access to popular RNA-Seq tools, within python, via easy-to-use high level APIs. Pipelines can be customized by integrating new python code, third-party programs, or python libraries. Users can create checkpoints in the pipeline or integrate **`pyrpipe`** into a workflow management system, thus allowing execution on multiple computing environments. **`pyrpipe`** produces detailed analysis, and benchmark reports which can be shared or included in publications. **`pyrpipe`** is implemented in python and is compatible with python versions 3.6 and higher. To illustrate the rich functionality of **`pyrpipe`**, we provide case studies using RNA-Seq data from GTEx, SARS-CoV-2-infected human cells, and Zea mays. All source code is freely available at https://github.com/urmi-21/pyrpipe; the package can be installed from the source or from PyPI (https://pypi.org/project/pyrpipe). Documentation is available at (http://pyrpipe.rtfd.io).

## INTRODUCTION

Since its inception, RNA-Seq has become the most widely used method to quantify transcript levels (1). A researcher can leverage the now-massive RNA-Seq data, encompassing multiple species, organs, genotypes, and conditions (2). Integrated analysis of aggregations of these diverse RNA-Seq samples enables exploration of changes in gene expression over time and across different biological conditions (3).

A major challenge in bioinformatics analysis of RNA-Seq datasets is implementing data processing pipelines in an efficient, modular, and reproducible manner (4, 5, 6, 7). A majority of existing bioinformatics tools are standalone linux programs, executed via the shell; writing bioinformatic pipelines as shell, perl, or python scripts is a very common practice among bioinformaticians. Scripting, although powerful and flexible, can be difficult to develop, understand, maintain, and debug, especially for complex workflows.

Since most bioinformatics tools are available as commands executed via a command-line interface (CLI), they must be specified inside a bash or a other scripting language for automated execution. The resulting scripts can often contain significant *boilerplate code* as user repeat the commands and parameters in the script. Managing the tools’ parameters, and making changes become difficult and error prone for complex pipelines. Controlling these parameters are essential for reproduciblity but are often not included in the methods and can be hard to track if not well documented. Moreover, no easy to use framework exists to define tools and parameters and modify them them at runtime.

Here we present `pyrpipe`, a lightweight python package for users to code and execute computational pipelines in an object oriented manner, in pure python. No new *workflow* syntax that is specific to `pyrpipe` is required. `pyrpipe` delivers an easy-to-use framework to easily import any Unix executable command or third-party tool as reusable python objects.

`pyrpipe` provides many useful features such as extensive logging and reports, loading tool options from yaml files, dry-run mode to check dependencies and targets, resuming of interrupted of jobs and saving `pyrpipe` sessions. We have incorporated APIs to popular RNA-Seq tools in `pyrpipe` to enable easy RNA-seq processing – from downloading raw data to trimming, alignment and assembly or quantification.

`pyrpipe` is designed to be easily integrated into a workflow management system, including the popular Snakemake (6), NextFlow (7) or Toil (8). The workflow management system then can scale and manage jobs on clusters and schedule independent jobs for parallel processing, facilitating scaleable pipelines and thus enabling analysis of large amounts of RNA-Seq data. Meanwhile, `pyrpipe` facilitates ease-of-implementation, reproducibility, understandibility, and modification of the pipeline.

## MATERIALS AND METHODS

### Overview

We developed `pyrpipe` to provide a light-weight and easy-to-use python framework for implementing bioinformatics or other computational analysis pipelines. The `pyrpipe` framework include: 1. high level APIs to popular RNA-Seq tools; 2. a general API to import any Unix command/tool into python, enabling use of any bioinformatics tool; and 3. extensive monitoring and logging details of the commands executed.

Thus, `pyrpipe` allows users to import any Unix executable command/tool into the python ecosystem and implement pipelines in pure-python incorporating their own python code, existing python libraries and third-party programs. `pyrpipe` is packaged as a python library and can be installed via PyPI or conda. An advantage of using the python platform is that it is widely used, free, flexible, object-oriented, has high-level data structures (9, 10), and a growing repository of > 200,000 packages and tools.

### The pyrpipe framework

`pyrpipe` enables users to code pipelines in an object oriented manner, using specialized API classes provided by `pyrpipe`. Using classes to encapsulate the various “tools” and “data” is the key structural concept of `pyrpipe`. Based on the principle of abstraction, `pyrpipe` hides the unnecessary details and provides the user with simple-to-use objects and functions. Thus, the “tools” can be easily accessed as objects in a re-usable manner, while ensuring that associated data and parameters are consistently accessible within that object and retaining the full functionality of the command/tool.

At its core, `pyrpipe` implements the “Runnable” class, which contains all necessary methods for *importing* any executable command/tool in python. Users may simply create an instance of the “Runnable” class to import any Unix tool in python, or can create a new class extending the “Runnable” class with additional specialized functionalities. Users can employ several helper functions defined in the modules *pyrpipe_engine*, and *`pyrpipe`.utils* to access frequently used operations.

Upon creation of a “Runnable” object, the tool options and arguments, if specified in a yaml file, are automatically loaded and stored in the object. These can be dynamically modified during execution. Parameter loading is robust and can ignore incorrect or misspelled options raising a warning.

The “run” method, implemented in the “Runnable” class, is responsible for executing the commands. The “run” method allows users to specify required dependencies and target files. The required files, if not present, will cause an exception. The target (output) files are equivalent to targets in GNU make; execution is skipped if the target files already exist. If a workflow is interrupted during execution it is resumed from where the last incomplete step, unless forced by the user to re-run from the beginning. Target files for interrupted runs are automatically removed. The “run” method integrates the command and its options and passes them to the `pyrpipe`_engine module, where the commands are executed and extensively logged.

### APIs for RNA-Seq processing

`pyrpipe` provides high-level APIs, to access full functionality of 11 popular RNA-Seq analysis tools that expedite and enhance implementation of RNA-Seq pipelines that can be readily shared, modified, or reused, including a dedicated module to easily access and manage RNA-Seq data available from the National Center for Biotechnology Research Sequence Read Archives (NCBI-SRA) database (2). These API classes are implemented inside several highly cohesive modules: (*sra, mapping, alignment, quant, qc, tools*). Each module has been designed to capture steps integral to RNA-Seq analysis: 1) access NCBI-SRA and manage raw RNA-Seq data; 2) quality control; 3) read alignment; 4) transcript assembly; and 5) transcript quantification. (Supplementary Table 1 and Supplementary Figures 1, 2). We have built and integrated these APIs into the `pyrpipe` package, such that any RNA-Seq processing pipeline can be intuitively executed by the researcher while writing minimal code (Figure 1).

**Figure 1.**
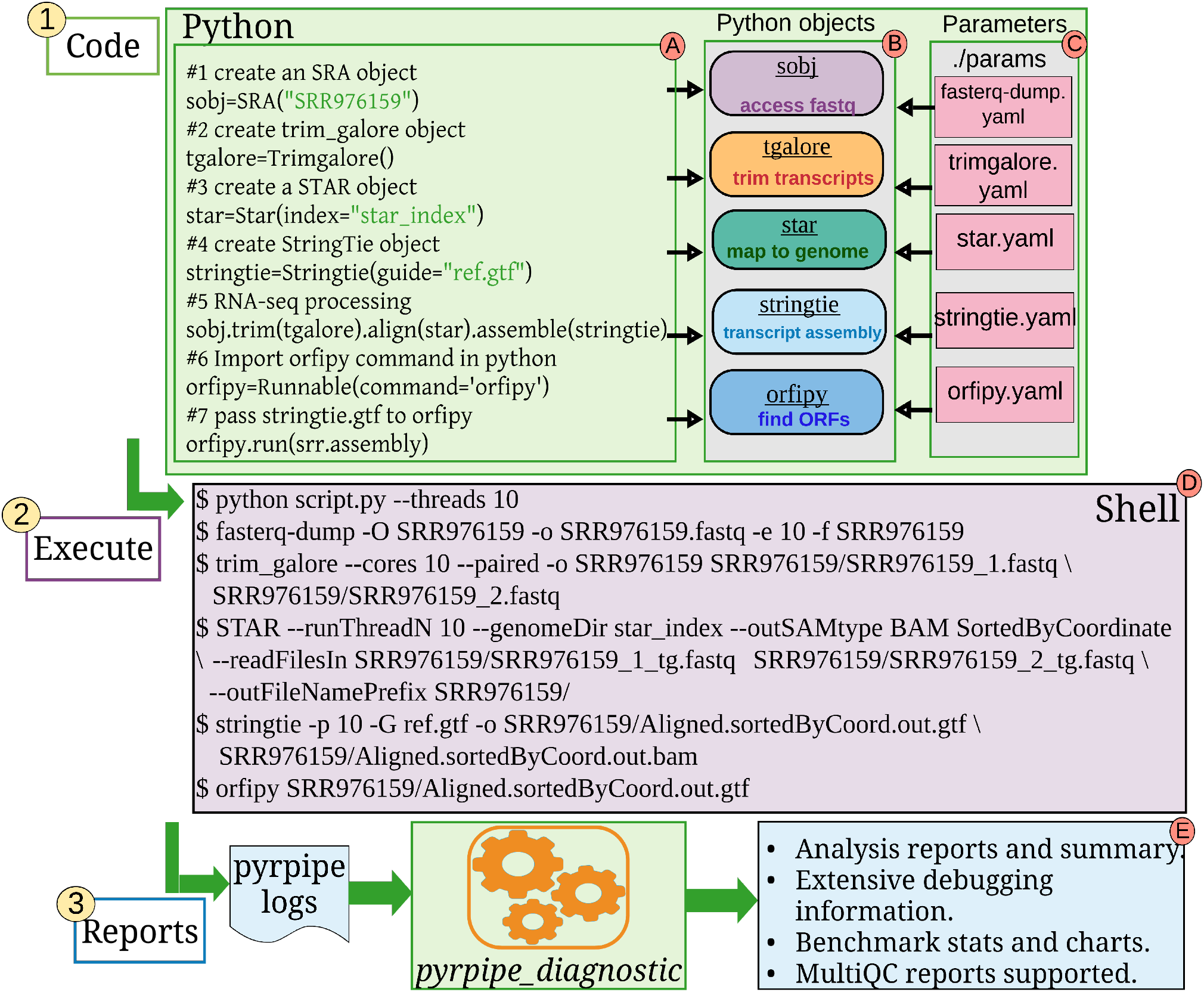
A simple example demonstrating `pyrpipe`. `pyrpipe` is represented by green boxes. The user writes the code in python and creates python *objects* of specific `pyrpipe` classes that provide API to RNA-Seq tools (Box A). Each *object* and encapsulates specific *methods* and *data* (Box B). Upon creation of each object, parameters specified in *yaml* files are loaded in the object (Box C). During execution, shell commands are automatically constructed and executed by the `pyrpipe` APIs (Box D). After execution, the *pyrpipe_diagnostic* tool generates the data analyses and diagnostic reports from the logs (Box E).

### Flexibility in pipeline execution, debugging, and pipeline sharing

`pyrpipe` flexibility extends to enabling the user to choose how to execute and handle exceptions and errors to modify their pipeline’s behavior.

Users can create checkpoints in the pipeline, save the current `pyrpipe` *session*, and resume later. This is particularly useful for running different blocks of a workflow in different environments. For example, on a typical high performance computing (HPC) cluster, a researcher might use a dedicated data-transfer node to retrieve data from SRA and then use compute nodes for data processing.

`pyrpipe` allow users to *dry run* the pipeline. During a *dry run*, all the dependencies and targets are checked and any missing dependencies or existing targets are reported to the user. No commands are executed via the *pyrpipe_engine*. Users can examine the *dry run* output to verify the commands and options.

`pyrpipe`’s logging features enable efficient error detection and reports (Fig. 1). Errors and extensive environment information, such as operating system and python version, along with version and path information for each program used within the pipeline, are all logged. `pyrpipe` logs are saved in JavaScript Object Notation (JSON) format for easy parsing by `pyrpipe` and other software (Supplementary Table 2).

The *pyrpipe_vdiagnostic* command can be invoked to generate comprehensive reports about the analysis, benchmark comparisons (Supplementary Figure 5), shell scripts and MultiQC reports (11). These reports, along with the python scripts, can be shared or included with publications to ensure reproducibility.

The default pyrpipe behaviour for logging, dry-run, and reports, can be easily modified by supplying pyrpipe with specific options via command-line or by specifying these in a *pyrpipe_conf.yaml* file.

### Reproducible analysis

Reproducibility can be a major challenge in bioinformatics studies because of heavy computational intensive tasks that depend on a number of software and system libraries. Reproducibility can be ensured by controlling execution environments via environment managers such as Anaconda, container systems such a Docker, or isolated virtual machines (4).

`pyrpipe` is a python package available through bioconda (12) and can be easily installed and managed within conda environments, containers or VMs. We have included in `pyrpipe` documentation the recommended way of installing the required tools for RNA-Seq analysis via bioconda.

Besides the user controlling the execution environment, `pyrpipe` adds several layers to enhance reproducibility of analysis. `pyrpipe` creates a local copy of the pipeline script so that user has access to the exact pipeline code later. `pyrpipe` logs the MD5 checksums of the pipeline script and any input files provided as arguments. Thus, the user can verify which scripts and input files were used in the analysis. We recommend users to use a version control software such as Git to keep a track of the changes to the scripts.

`pyrpipe` allows and encourages users to define separate yaml files for the tool parameters. This makes it easy to modify, manage, share and reproduce computational analysis on different data and platforms. Further, `pyrpipe` logs contain detailed information about all the tools/commands used and their versions, which can be utilized to re-build the environments.

## RESULTS

We evaluated `pyrpipe` by three case studies, each illustrating a different aspect of what the tool can accomplish and how new functionality can be added.

### Case study 1: Scaling up pyrpipe to process 17,328 RNA-Seq samples from non-diseased human tissues

This case study demonstrates the ability of `pyrpipe` to process large amounts of data – 17,328 human RNA-Seq samples from the Genotype-Tissue Expression (GTEx V8) (13). We developed and implemented our pipeline to identify expressed human orphan genes in diverse tissues using `pyrpipe` (Supplementary Figure 3). This pipeline was run on the PSC Bridges (https://www.psc.edu/resources/bridges/) HPC system. The pipeline was scaled to run multiple batches of RNA-Seq samples in parallel on multiple nodes. Code for this project is available https://github.com/urmi-21/pyrpipe/tree/master/case_studies/GTEx_processing.

To assess the results of our pipeline, we compare the median TPMs of annotated genes for two types of adipose tissue, as processed by `pyrpipe` and by the GTEx portal (Figure 2).

**Figure 2.**
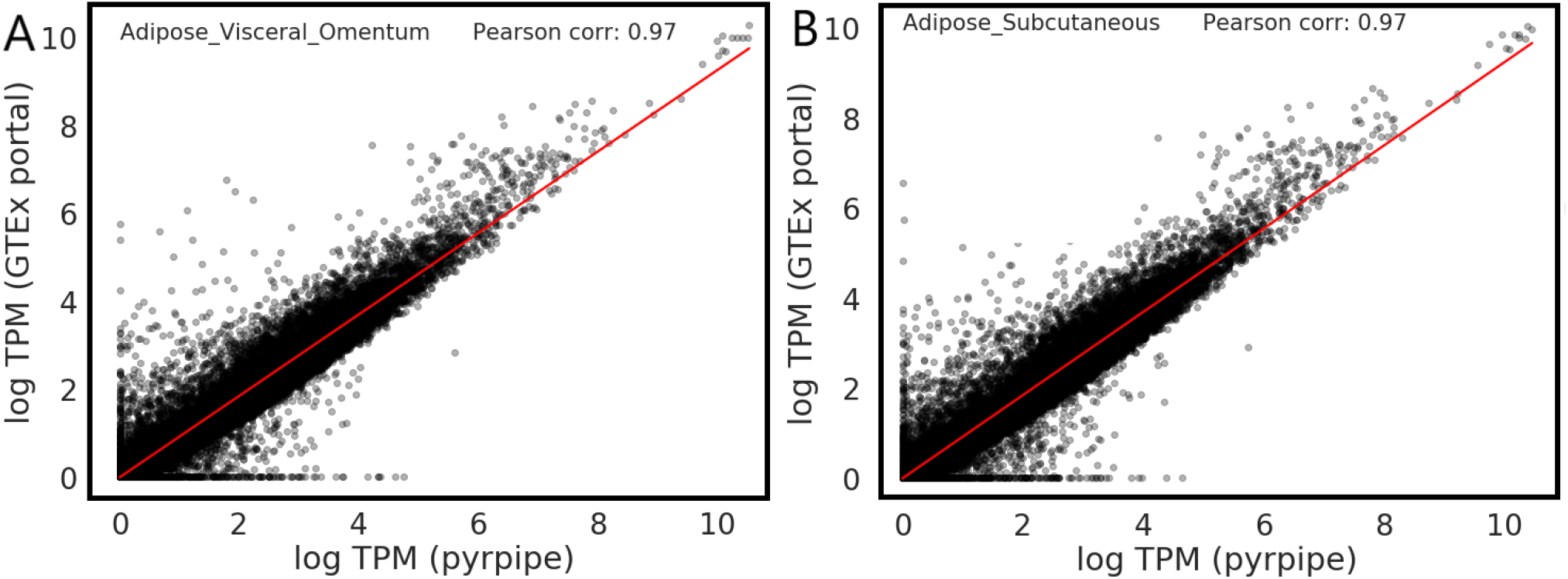
Comparison of median TPMs for two tissue types **A.** Visceral Adipose and **B.** Subcutaneous Adipose. Y-axis shows the logged Median TPMs computed by the GTEx portal pipeline. X-axis shows the logged Median TPMs computed by our pipeline implemented with `pyrpipe`. Pearson correlations are 0.97%. Differences in quantification of several 100 genes are likely due differences in reference annotations.

### Case study 2: Integrating pyrpipe within a workflow manager to quantify gene expression in COVID-19 samples for exploratory analysis

We implemented `pyrpipe` within two workflow management systems, Snakemake (6) and NextFlow (7), selecting these specifically because they are widely used by the bioinformatics community. Snakemake and NextFlow were independently used to implement, manage and execute the pipeline for multiple RNA-Seq samples in parallel on a single cluster. This pipeline quantifies RNA-Seq data from a COVID-19 study (14), and provides output that can be directly analyzed by biologists, using the versatile Java software for exploratory analysis of large datasets, MetaOmGraph (MOG) (15).

Specifically, we use `pyrpipe` to download 29 RNA-Seq samples from NCBI-SRA published under the accession SRP287810, and quantify expression of annotated transcripts using salmon’s selective alignment approach (16, 17). The final transcript and gene level TPMs from each sample are merged into a single file to create a MetaOmGraph (3) project (*MOGproject-monocytes-60241genes-45samples-HCQtreat-2021-1-17*) for exploratory data analysis.

The code, data, and MOG project are available at https://github.com/urmi-21/pyrpipe/tree/master/case_studies/Covid_RNA-Seq.

### Case study 3: Use of pyrpipe for *de novo* transcriptome assembly

We used a new, high-quality genome of *Zea mays* B73 cultivar (19) as reference genome, and gathered RNA-Seq data from ten diverse samples (B73 cultivar), representing different tissue and development stages, for *de novo* transcriptome assembly (Supplementary Figure 4). Our pipeline identified a total of 57,916 distinct transcripts. Of these, 38,881 transcripts were homologous to UniProt proteins (20, 21). These transcripts could be non-coding RNAs (ncRNAs), “noise”, pseudogenes, or genes encoding unannotated, conserved proteins. The remaining 6,306 transcripts, with no similarity to any protein in the database, could potentially be ncRNAs, “noise”, or species-specific (“orphan”) genes (22). The transcript length and GC content distribution for transcripts with conserved CDS and transcripts are shown in Figure 3. The mean length of non-homologous transcripts (1,290 nt) is shorter than conserved transcripts (1,981 nt); mean GC content is indistinguishable (50.8% vs 50.6%). The median expression of non-homologous transcripts across the 10 RNA-Seq samples analyzed is lower than the median expression of conserved transcripts; however, in each sample, hundreds of non-homologous transcripts are more highly expressed than the mean of the conserved genes. These characteristics follow the same trend as those of conserved and orphan genes in the well-characterized *Arabidopsis thaliana* genome (18).

**Figure 3.**
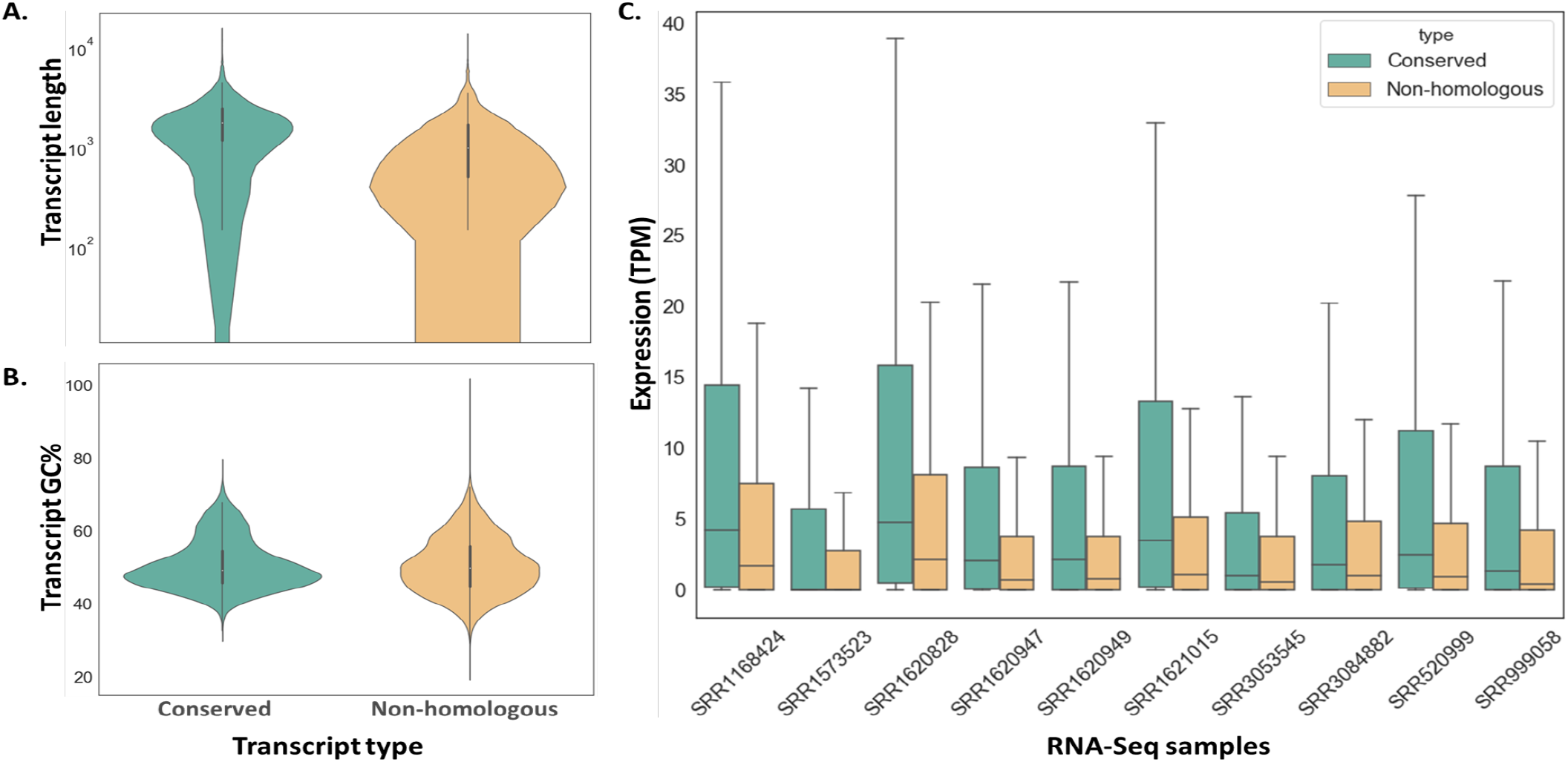
Comparison of length, %GC content, and expression level between conserved and non-homologous transcripts as identified by our pipeline. The length of non-homologous transcripts (mean = 1,290 nt) is shorter than conserved transcripts (mean = 1,981 nt). In contrast, mean GC content is indistinguishable between conserved and non-homologous transcripts (50.8% and 50.6%). The median expression of non-homologous transcripts across the 10 RNA-Seq samples analyzed is lower than the median expression of conserved transcripts. However, in each sample some non-homologous transcripts are more highly expressed than the mean of the conserved genes. These characteristic follow the same trend as those of conserved and orphan genes in the well-characterized *Arabidopsis thaliana* genome (18).

Further, median expression of these non-homologous transcripts across the RNA-Seq samples analyzed is lower than the median expression of conserved transcripts (Figure 3). Pipeline scripts, downstream analysis code and data are available at https://github.com/lijing28101/maize_pyrpipe.

### Comparison of pyrpipe to existing python libraries that can be used for RNA-Seq analysis

Several python libraries enable workflows to be specified. However, they do not provide a dedicated API suite for RNA-Seq data analysis. Instead, these frameworks depend on the user to explicitly write the commands and provide data, diminishing the reproducibility of the results.

We compared `pyrpipe` with two such python libraries that allow specifying bioinformatics pipeline - Ruffus (23) and Pypiper (http://code.databio.org/pypiper/). Ruffus is a python library for specifying and executing workflows. Ruffus allows users to specify pipeline tasks using several *“decorator”* functions. Pypiper is a python package for coding pipelines in python. It provides the “PipelineManager” class which a user can employ to execute commands in a serial manner. Pypiper has a built-in toolkit, NGSTk, to allow users to generate commonly used bioinformatics shell commands. These functions return commands as *string* objects that can be passed to “PipelineManager” for execution. Table 1 compares `pyrpipe` features with Ruffus and Pypiper.

**Table 1.** Comparison of `pyrpipe` features with Ruffus and Pypiper.

| Feature | `pyrpipe` | Ruffus | Pypiper |
| --- | --- | --- | --- |
| Latest update | 2021 | 2020 | 2019 |
| API to RNA-Seq tools | ✓ | ✗ | ✗ |
| Import tools as objects | ✓ | ✗ | ✗ |
| Auto-load tool parameters | ✓ | ✗ | ✗ |
| Dry run mode | ✓ | ✗ | ✗ |
| Resume Interrupted | ✓ | ✓ | ✓ |
| Exception handling | ✓ | ✓ | ✓ |
| Parallel execution support | ✗ | ✓ | ✗ |
| Logs/Reports | ✓/✓ | ✓/✗ | ✓/✓ |

## DISCUSSION AND CONCLUSION

The `pyrpipe` package allows users to code and implement RNA-Seq workflows in an object oriented manner, purely using python. APIs to RNA-Seq tools make it straightforward to code RNA-Seq processing pipelines. Access to NCBI-SRA is automated, such that users can readily retrieve raw read RNA-Seq data. The downloaded raw RNA-Seq data and data files are automatically managed, and consistently accessed through *SRA* objects. Users need not keep track of data files or paths, as these are integrated with `pyrpipe` objects. `pyrpipe` workflows can be modified using python’s control flow abilities and a user can create complex, reproducible, workflow structures. Any third party tool, Unix command, or script can be integrated into `pyrpipe` for additional data processing capability. `pyrpipe` logs and reports enable debugging and reproducibility.

Analysis of the 17,328 GTEx RNA-Seq samples was easily scaled using `pyrpipe` alone, by creating smaller *batches* of samples and submitting the processing jobs in parallel, on an HPC system with a slurm job scheduler.

When building more complex and scalable workflows, it may be more efficient to integrate `pyrpipe` into a workflow management system. This can easily be done, as shown in our second case study. Workflow management systems are developed for robust and easy implementation of computational pipelines; nevertheless, they differ significantly in terms of workflows, definitions, job scheduling, and features (5, 6, 7, 8). For example, Snakemake uses a “pull-based” strategy to check for specific output files and schedule jobs accordingly (5, 6), whereas Nextflow uses a “push-based” scheme in which a “process” defined in the workflow pushes its outputs to downstream “processes” (7, 24). The SciPipe (5) workflow library is written in the GO language; similar to Nextflow it implements dataflow based task scheduling. Toil (8) provides explicit application programming interfaces (APIs) for defining static or dynamic tasks and supports common workflow language (CWL) and multiple cloud environments (8, 25). Hence, users need to make informed decisions if choosing a workflow management system (26) for `pyrpipe`.

The modular design of `pyrpipe` permits users to write *py?honic* code, which is easy to read, manage, and share. Because of the rapid emergence of new bioinformatics tools, this design feature is particularly important. From a developer’s perspective, `pyrpipe`’s modularity facilitates reuse and extensibility; new tools/APIs can be easily integrated into `pyrpipe` and promotes sustainability.

`pyrpipe` will appeal to users who are looking for simple, fast way to deploy small or large scale RNA-Seq processing pipelines. Straightforward implementation, seamless integration, execution and sharing of RNA-Seq workflows makes it an ideal choice for users with less computational expertise.

## DATA AVAILABILITY

We subscribe to FAIR data and software practices (27). `pyrpipe` source code is available at https://github.com/urmi-21/pyrpipe. `pyrpipe` source code (v0.0.5) can be accessed via DOI: 10.5281/zenodo.4448373. The `pyrpipe` package can be installed from the source, from PyPi (https://pypi.org/project/pyrpipe) or from bioconda (https://anaconda.org/bioconda/pyrpipe). Extensive documentation to guide users on how to use `pyrpipe` and the APIs implemented within it is available on Read the Docs (http://pyrpipe.rtfd.io).

## SUPPLEMENTARY DATA

Supplementary Data are available at https://github.com/urmi-21/pyrpipe.

## REFERENCES

1. Stark, R., Grzelak, M., and Hadfield, J. (2019) RNA sequencing: the teenage years. Nature Reviews Genetics, 20(11), 631–656.

2. Kodama, Y., Shumway, M., and Leinonen, R. (2011) The Sequence Read Archive: explosive growth of sequencing data. Nucleic acids research, 40(D1), D54–D56.

3. Singh, U., Hur, M., Dorman, K., and Wurtele, E. S. (01, 2020) MetaOmGraph: a workbench for interactive exploratory data analysis of large expression datasets. Nucleic Acids Research, gkz1209.

4. Grüning, B., Chilton, J., Köster, J., Dale, R., Soranzo, N., van den Beek, M., Goecks, J., Backofen, R., Nekrutenko, A., and Taylor, J. (2018) Practical computational reproducibility in the life sciences. Cell systems, 6(6), 631–635.

5. Lampa, S., Dahlö, M., Alvarsson, J., and Spjuth, O. (2019) SciPipe: A workflow library for agile development of complex and dynamic bioinformatics pipelines. GigaScience, 8(5), giz044.

6. Köster, J. and Rahmann, S. (2012) Snakemake—a scalable bioinformatics workflow engine. Bioinformatics, 28(19), 2520–2522.

7. Di Tommaso, P., Chatzou, M., Floden, E. W., Barja, P. P., Palumbo, E., and Notredame, C. (2017) Nextflow enables reproducible computational workflows. Nature biotechnology, 35(4), 316.

8. Vivian, J., Rao, A. A., Nothaft, F. A., Ketchum, C., Armstrong, J., Novak, A., Pfeil, J., Narkizian, J., Deran, A. D., Musselman-Brown, A., et al. (2017) Toil enables reproducible, open source, big biomedical data analyses. Nature biotechnology, 35(4), 314.

9. Kossaifi, J., Panagakis, Y., Anandkumar, A., and Pantic, M. (2019) Tensorly: Tensor learning in python. The Journal of Machine Learning Research, 20(1), 925–930.

10. Kanterakis, A., Iatraki, G., Pityanou, K., Koumakis, L., Kanakaris, N., Karacapilidis, N., and Potamias, G. (2019) Towards reproducible bioinformatics: the OpenBio-C scientific workflow environment. In 2019 IEEE 19th International Conference on Bioinformatics and Bioengineering (BIBE) IEEE Computer Society pp. 221–226.

11. Ewels, P., Magnusson, M., Lundin, S., and Käller, M. (2016) MultiQC: summarize analysis results for multiple tools and samples in a single report. Bioinformatics, 32(19), 3047–3048.

12. Grüning, B., Dale, R., Sjödin, A., Chapman, B. A., Rowe, J., Tomkins-Tinch, C. H., Valieris, R., and Köster, J. (2018) Bioconda: sustainable and comprehensive software distribution for the life sciences. Nature methods, 15(7), 475–476.

13. Consortium, G. et al. (2017) Genetic effects on gene expression across human tissues. Nature, 550(7675), 204–213.

14. Rother, N., Yanginlar, C., Lindeboom, R. G., Bekkering, S., van Leent, M. M., Buijsers, B., Jonkman, I., de Graaf, M., Baltissen, M., Lamers, L. A., et al. (2020) Hydroxychloroquine inhibits trained immunity-implications for COVID-19. medRxiv,.

15. Singh, U., Hur, M., Dorman, K., and Wurtele, E. S. (2020) MetaOmGraph: a workbench for interactive exploratory data analysis of large expression datasets. Nucleic Acids Research, 48(4), e23–e23.

16. Patro, R., Duggal, G., Love, M. I., Irizarry, R. A., and Kingsford, C. (2017) Salmon provides fast and bias-aware quantification of transcript expression. Nature methods, 14(4), 417.

17. Srivastava, A., Malik, L., Sarkar, H., Zakeri, M., Almodaresi, F., Soneson, C., Love, M. I., Kingsford, C., and Patro, R. (2020) Alignment and mapping methodology influence transcript abundance estimation. Genome biology, 21(1), 1–29.

18. Arendsee, Z. W., Li, L., and Wurtele, E. S. (2014) Coming of age: orphan genes in plants. Trends in plant science, 19(11), 698–708.

19. Hufford, M. B., Seetharam, Arun S, Woodhouse, M. R., Chougule, K. M., Ou, S., Liu, J., Ricci, W. A., Guo, T., Olson, A., Qiu, Y., Della Coletta, R., Tittes, S., Hudson, A. I., Marand, A. P., Wei, S., Lu, Z., Wang, B., Tello-Ruiz, M. K., Piri, R. D., Wang, N., Kim, D. w., Zeng, Y., O’Connor, C. H., Li, X., Gilbert, A. M., Baggs, E., Krasileva, K. V., Portwood, J. L., Cannon, E. K., Andorf, C. M., Manchanda, N., Snodgrass, S. J., Hufnagel, D. E., Jiang, Q., Pedersen, S., Syring, M. L., Kudrna, D. A., Llaca, V., Fengler, K., Schmitz, R. J., Ross-Ibarra, J., Yu, J., Gent, J. I., Hirsch, C. N., Ware, D., and Dawe, R. K. (2021) De novo assembly, annotation, and comparative analysis of 26 diverse maize genomes. bioRxiv,.

20. Consortium, U. (2019) UniProt: a worldwide hub of protein knowledge. Nucleic acids research, 47(D1), D506–D515.

21. Altschul, S. F., Gish, W., Miller, W., Myers, E. W., and Lipman, D. J. (1990) Basic local alignment search tool. Journal of molecular biology, 215(3), 403–410.

22. Singh, U. and Wurtele, E. S. (2020) Genetic Novelty: How new genes are born. Elife, 9, e55136.

23. Goodstadt, L. (2010) Ruffus: a lightweight Python library for computational pipelines. Bioinformatics, 26(21), 2778–2779.

24. Strozzi, F., Janssen, R., Wurmus, R., Crusoe, M. R., Githinji, G., Di Tommaso, P., Belhachemi, D., Möller, S., Smant, G., de Ligt, J., et al. (2019) Scalable workflows and reproducible data analysis for genomics. In Evolutionary Genomics pp. 723–745 Springer.

25. Leipzig, J. (2017) A review of bioinformatic pipeline frameworks. Briefings in bioinformatics, 18(3), 530–536.

26. Jackson, M., Wallace, E., and Kavoussanakis, K. (2020) Using rapid prototyping to choose a bioinformatics workflow management system. bioRxiv,.

27. Wilkinson, M. D., Dumontier, M., Aalbersberg, I. J., Appleton, G., Axton, M., Baak, A., Blomberg, N., Boiten, J.-W., da Silva Santos, L. B., Bourne, P. E., et al. (2016) The FAIR Guiding Principles for scientific data management and stewardship. Scientific data, 3.

