## Supplementary Data for "pyrpipe: a python package for RNA-Seq workflows"

### orfipy: a fast and flexible tool for extracting ORFs (Supplementary Materials)

#### Supplementary Information

| Module name | Class name | API for | Purpose |
|---|---|---|---|
| sra | SRA | sra-tools[1] | Access NCBI-SRA database and manage Fastq files |
| mapping | Hisat2 | Hisat2[2] | Read alignment |
| | Star | STAR[3] | Read alignment |
| | Bowtie2 | Bowtie2[4] | Read alignment |
| assembly | Stringtie | StringTie[5] | Transcript assembly |
| | Cufflinks | Cufflinks[6] | Transcript assembly |
| quant | Kallisto | Kallisto[7] | Transcript quantification |
| | Salmon | Salmon[8] | Transcript quantification |
| qc | Trimgalore | Trim Galore[9] | Quality control |
| | BBDuk | BBDuk[10] | Quality control |
| tools | Samtools | Samtools[11] | Processing read alignments |

**Supplementary Table 1.** Currently implemented *pyrpipe* modules. Each module contain multiple classes containing APIs for different RNA-Seq tools.

| Key | Description |
|---|---|
| cmd | shell command executed |
| starttime | Time at the start of execution |
| runtime | Total runtime |
| exitcode | The return code |
| stdout | stdout returned by the program |
| stderr | stderr returned by the program |
| objectid | Id of an object used with the command |
| commandname | Name of the command |
| python | Python version |
| os | Operating system |
| cpu | CPU information |
| syspath | Python's sys.path |
| sysmodules | Python's sys.modules |
| name | Name of the program executed |
| version | Version of the program |
| path | Path to the program on disk |

**Supplementary Table 2.** Table showing description of the JSON *keys* stored in `pyrpipe` logs

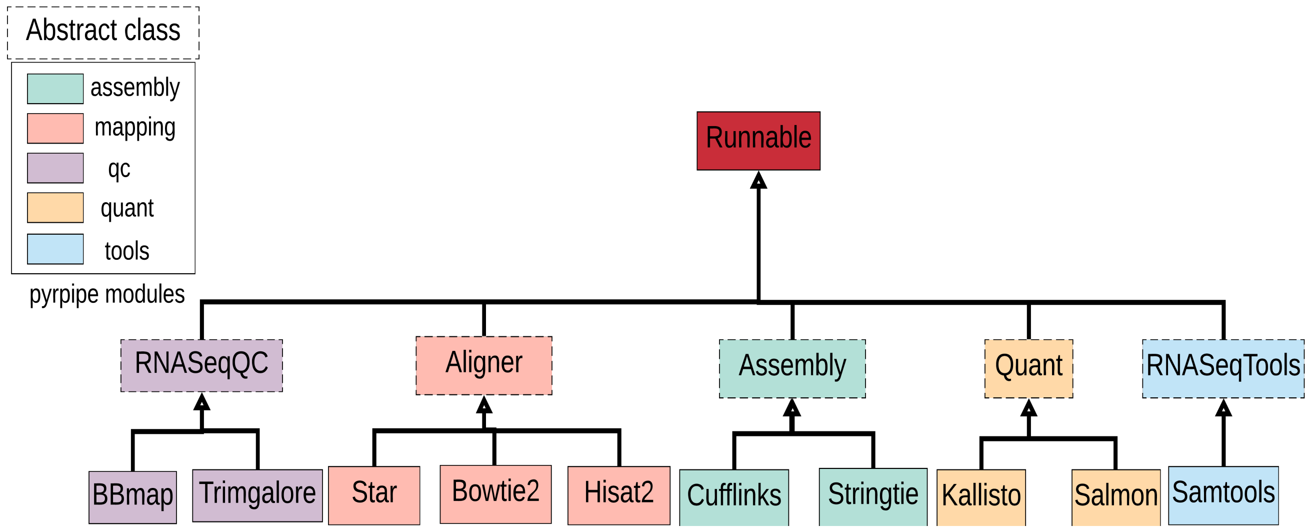


**Supplementary Figure 1.** A class diagram showing `pyrpipe's` class hierarchy. Classes in the same `pyrpipe` module share the same color; dotted rectangles represent abstract classes.

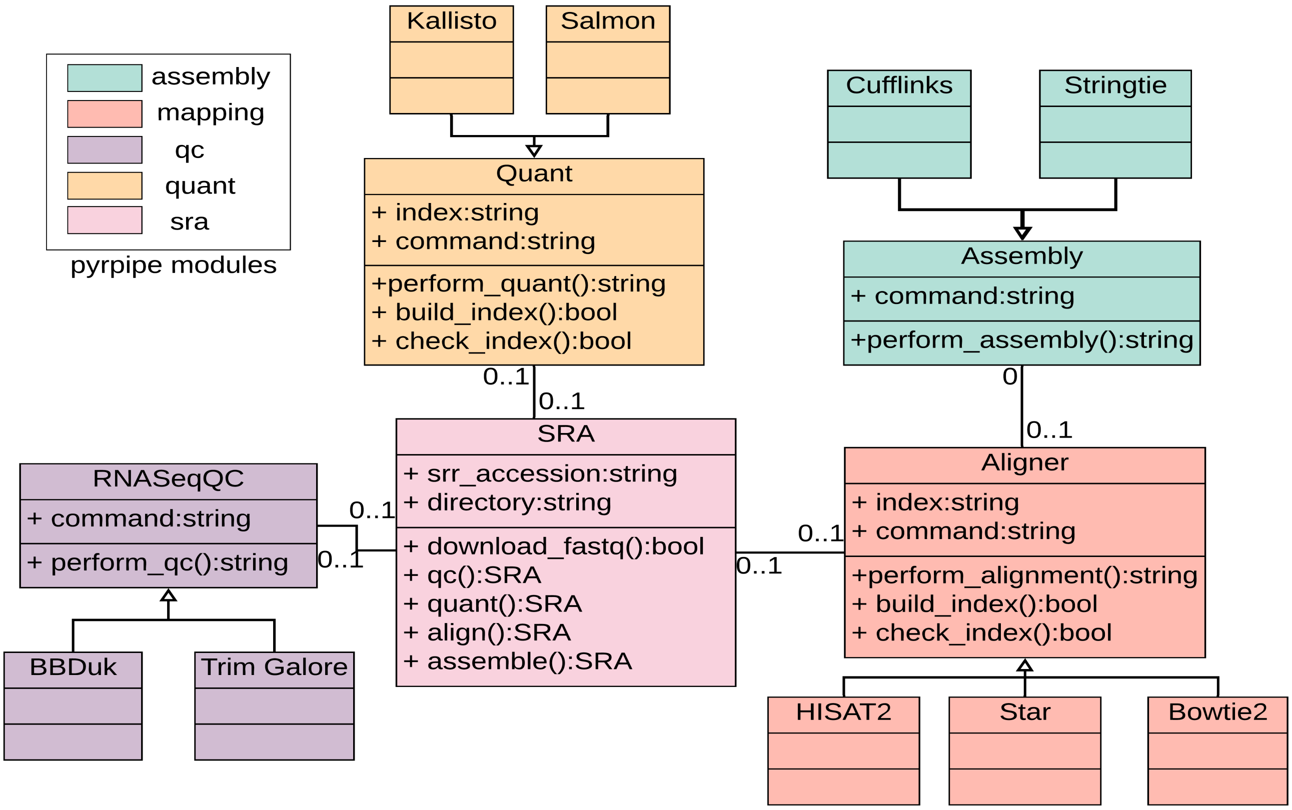


**Supplementary Figure 2.** A UML class diagram showing `pyrpipe's` RNA-Seq API classes and relationships among them. Classes in the same `pyrpipe` module share the same color.

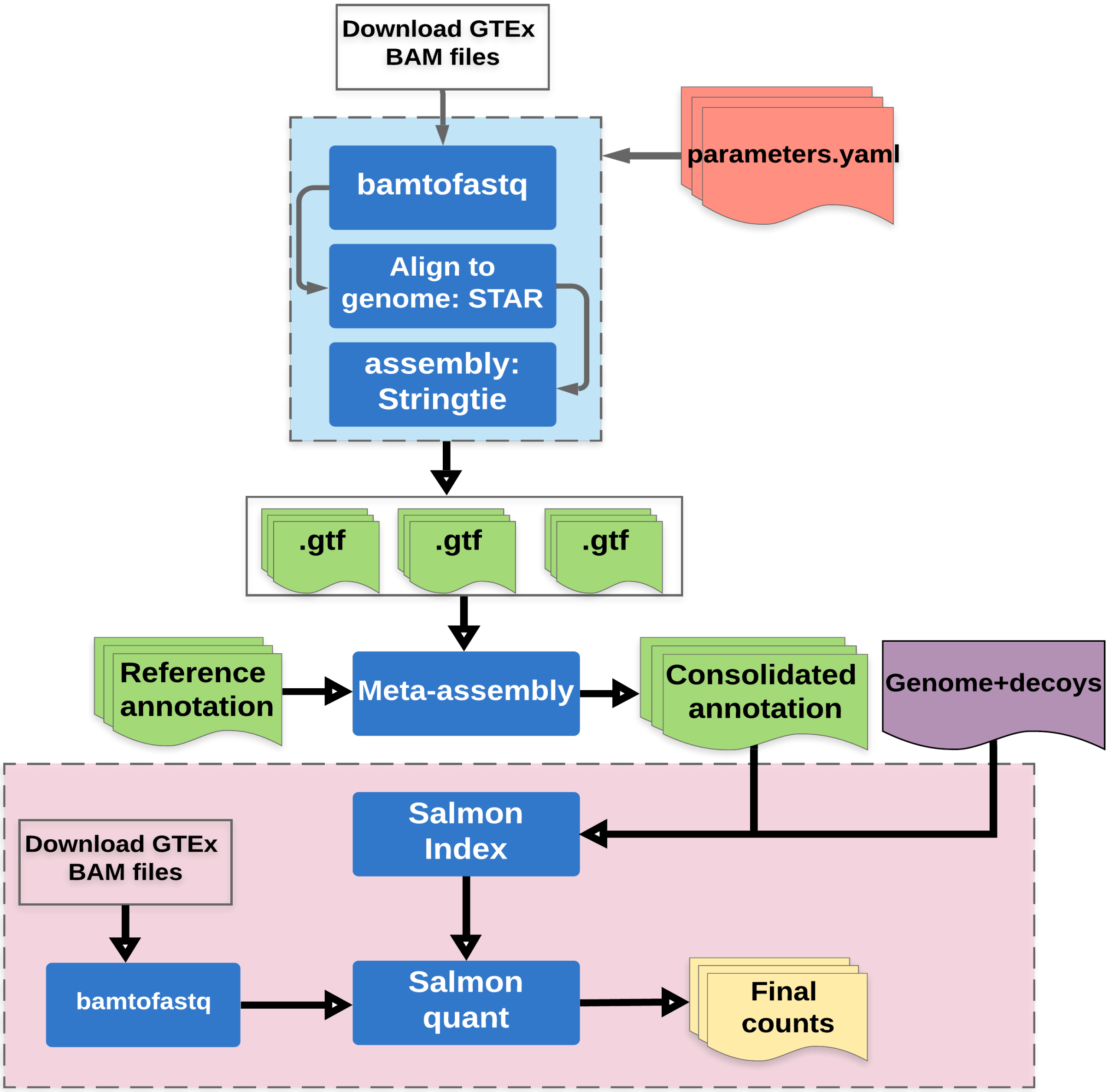


**Supplementary Figure 3.** A flowchart showing steps used to process GTEx RNA-Seq data. The Alignment and quantification pipelines are implemented in `pyrpipe`. For alignment, BAM files were downloaded from GTEx and converted to fastq using biobambam2[12]. STAR[3] was run in 2-pass alignment mode to align reads to the human reference genome. Stringtie[5] was used to assemble transcripts. Raw data files were deleted as after they were processed to save disk space. Individual transcriptomes were consolidated into single transcriptome using our unpublished meta-assembly pipeline which is similar to[13]. Salmon[8] index was build using human annotated and novel identified transcripts. Human whole genome sequence was used as a decoy. During quantification phase, BAM files were downloaded from GTEx, converted to fastq (using biobambam2) and passed to salmon quant for quantification.

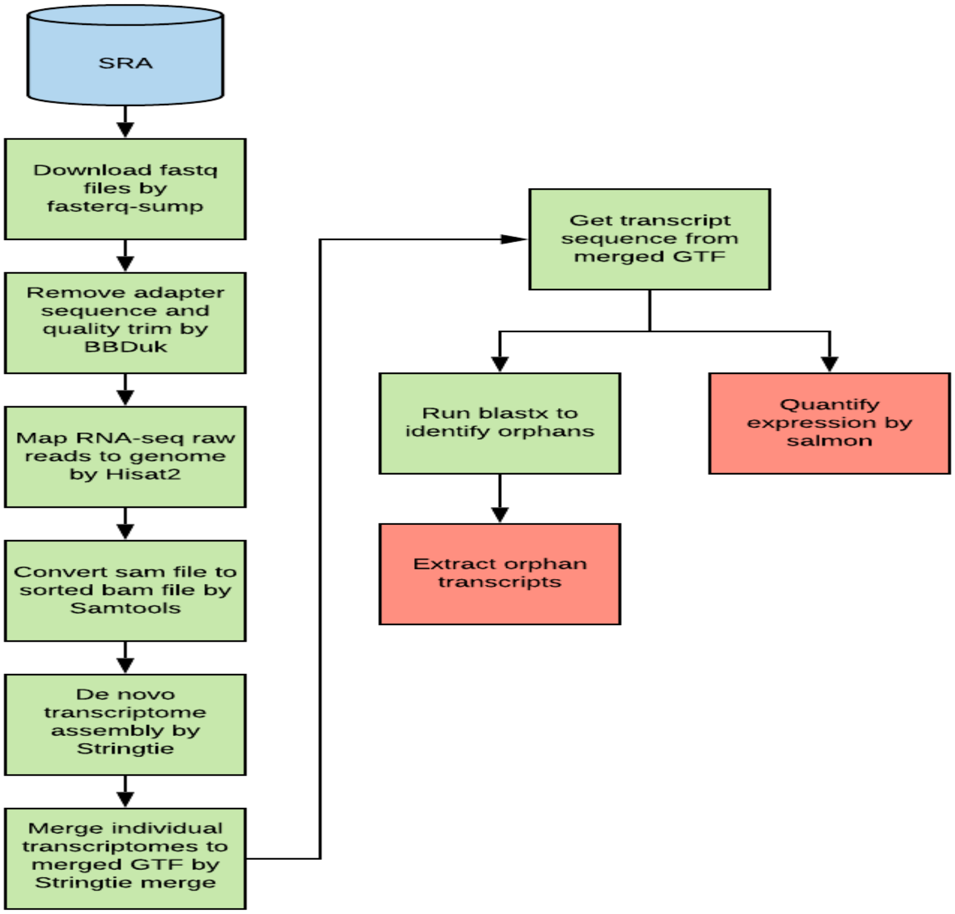


**Supplementary Figure 4.** A flowchart showing the pipeline implemented in `pyrpipe` to identify potentially orphan coding transcripts in *Zea mays*.

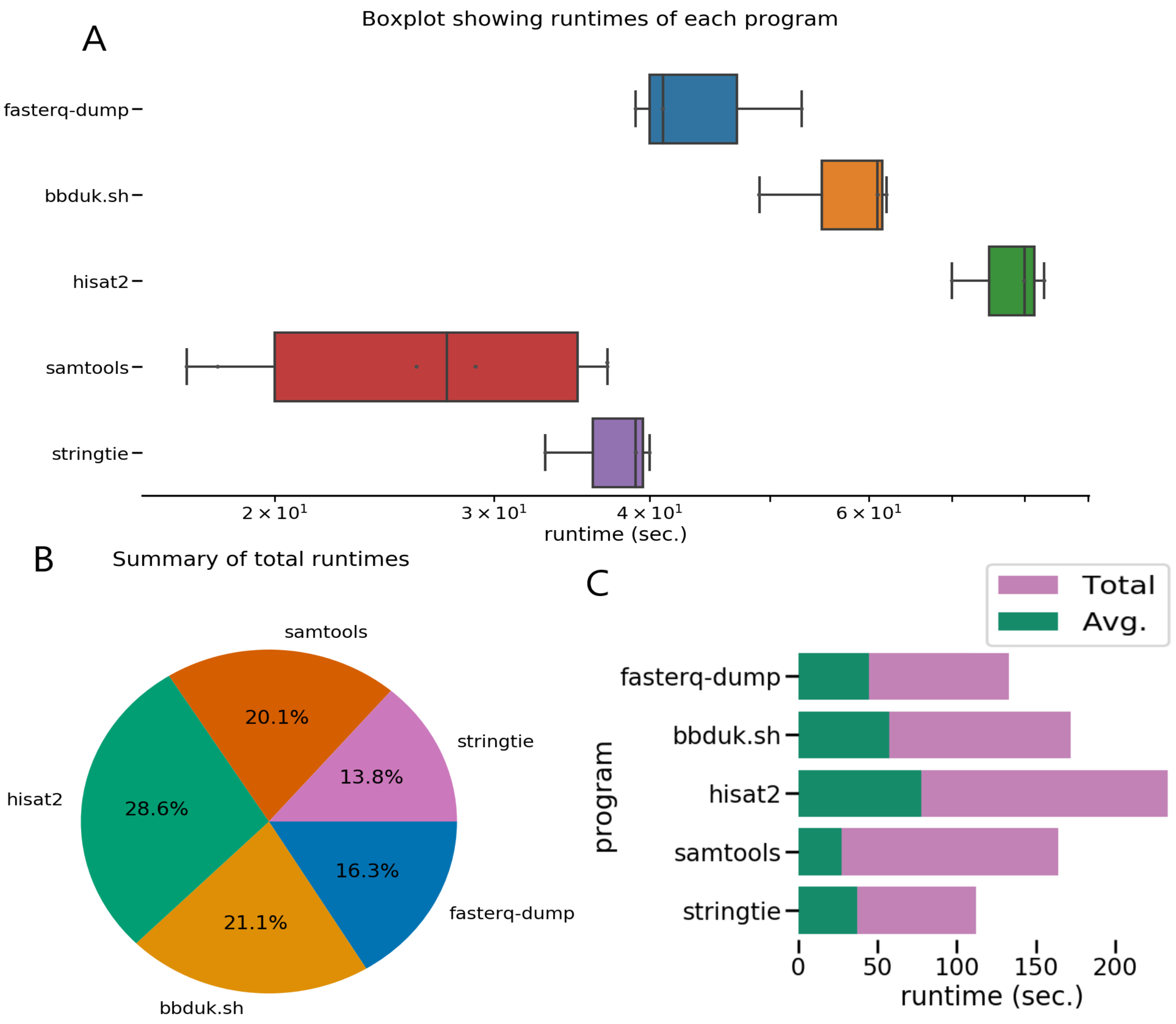


**Supplementary Figure 5.** Example of plots generated using pyrpipe_diagnostic benchmark command. **A.** Boxplots showing distribution of walltimes of different tools/commands used in the pipeline. **B.** A pie-chart showing how much each tool contributed to total runtime of the pipeline. **C.** Bar plot showing average and total runtimes of each tool tin the pipeline. The pipeline code and data is available from https://github.com/urmi-21/pyrpipe/tree/master/case_studies/Athaliana_transcript_assembly.

#### References


1. Sherry, S. & Xiao, C. Ncbi sra toolkit technology for next generation sequence data. In *Plant and Animal Genome XX Conference (January 14-18, 2012). Plant and Animal Genome* (2012).
2. Kim, D., Paggi, J. M., Park, C., Bennett, C. & Salzberg, S. L. Graph-based genome alignment and genotyping with hisat2 and hisat-genotype. *Nat. biotechnology* **37**, 907–915 (2019).
3. Dobin, A. *et al.* Star: ultrafast universal rna-seq aligner. *Bioinformatics* **29**, 15–21 (2013).
4. Langmead, B. & Salzberg, S. L. Fast gapped-read alignment with bowtie 2. *Nat. methods* **9**, 357 (2012).
5. Pertea, M. *et al.* Stringtie enables improved reconstruction of a transcriptome from rna-seq reads. *Nat. biotechnology* **33**, 290 (2015).
6. Trapnell, C. *et al.* Transcript assembly and quantification by rna-seq reveals unannotated transcripts and isoform switching during cell differentiation. *Nat. biotechnology* **28**, 511 (2010).
7. Bray, N. L., Pimentel, H., Melsted, P. & Pachter, L. Near-optimal probabilistic rna-seq quantification. *Nat. biotechnology* **34**, 525 (2016).
8. Patro, R., Duggal, G., Love, M. I., Irizarry, R. A. & Kingsford, C. Salmon provides fast and bias-aware quantification of transcript expression. *Nat. methods* **14**, 417 (2017).
9. Krueger, F. Trim galore. *A wrapper tool around Cutadapt FastQC to consistently apply quality adapter trimming to FastQ files* (2015).
10. Bushnell, B. Bbtools software package. *URL http://sourceforge. net/projects/bbmap* (2014).
11. Li, H. *et al.* The sequence alignment/map format and samtools. *Bioinformatics* **25**, 2078–2079 (2009).
12. Tischler, G. & Leonard, S. biobambam: tools for read pair collation based algorithms on bam files. *Source Code for Biol. Medicine* **9**, 13 (2014).
13. Seetharam, A. S. *et al.* Maximizing prediction of orphan genes in assembled genomes. *BioRxiv* (2019).
